# TF2Network: predicting transcription factor regulators and gene regulatory networks in Arabidopsis using publicly available binding site information

**DOI:** 10.1101/173559

**Authors:** Shubhada R. Kulkarni, Dries Vaneechoutte, Jan Van de Velde, Klaas Vandepoele

**Affiliations:** Ghent University, Department of Plant Biotechnology and Bioinformatics, (Technologiepark 927,) 9052 Ghent, Belgium; VIB Center for Plant Systems Biology, (Technologiepark 927,) 9052 Ghent, Belgium; Bioinformatics Institute Ghent, Ghent University, Technologiepark 927, 9052 Ghent, Belgium

## Abstract

A gene regulatory network (GRN) is a collection of regulatory interactions between transcription factors (TFs) and their target genes. GRNs control different biological processes and have been instrumental to understand the organization and complexity of gene regulation. Although various experimental methods have been used to map GRNs in *Arabidopsis thaliana*, their limited throughput combined with the large number of TFs makes that for many genes our knowledge about regulating TFs is incomplete. We introduce TF2Network, a tool that exploits the vast amount of TF binding site information and enables the delineation of GRNs by detecting potential regulators for a set of co-expressed or functionally related genes. Validation using two experimental benchmarks reveals that TF2Network predicts the correct regulator in 75-92% of the test sets. Furthermore, our tool is robust to noise in the input gene sets, has a low false discovery rate, and shows a better performance to recover correct regulators compared to other plant tools. TF2Network is accessible through a web interface where GRNs are interactively visualized and annotated with various types of experimental functional information. TF2Network was used to perform systematic functional and regulatory gene annotations, identifying new TFs involved in circadian rhythm and stress response.

## INTRODUCTION

Transcriptional regulation is one of the fundamental processes controlling gene expression and orchestrating gene activity resulting in phenotypic diversity. Key factors that drive this process are transcription factors (TFs) which regulate their target genes by recognizing short sequences on the DNA called TF binding sites (TFBSs). The full set of regulatory interactions between a TF and its target genes forms a gene regulatory network (GRN). GRNs are of major interest as they give an overview of how transcriptional regulation is organized at the genome-wide level. Although TFs are frequently classified as activators or repressors, gene regulation is typically controlled through the combinatorial control of different TFs, where context-dependency specifies how TFs modulate the expression of target genes (1,2). Based on the systematic analysis of TF chromatin immunoprecipitation (ChIP) experiments it has been estimated that in *Arabidopsis* a single gene can be regulated by up to 75 different TFs, highlighting the underlying network complexity (3,4).

Reliably identifying which TFs regulate which target genes is pivotal to understand how different biological processes like growth, development or stress response are transcriptionally controlled. Previous studies have started to study GRNs in specific cellular conditions using different experimental techniques. Whereas Brady and co-workers used a gene-centric yeast one hybrid (Y1H) approach to unravel the first tissue-specific GRN in Arabidopsis roots (5), Taylor-Teeples and co-workers used Y1H to unravel the GRN underlying secondary cell wall synthesis (6). Similarly, Y1H assays were used to generate a transcriptional network for root ground tissue and to identify new regulators for SHORTROOT-SCARECROW (7). In contrast to the *in vitro* Y1H method, ChIP-chip or ChIP-Seq are *in vivo* TF-centric techniques that determine TF binding and map the potential targets of an individual TF (3). Besides Y1H and ChIP, open chromatin profiling combines the TF footprint information obtained from DNAse I hypersensitivity (DH) assays with known TFBSs to build GRNs (8–10). This *in vivo* genome-wide approach is unbiased because it doesn’t need prior knowledge about potential TFs or target genes involved in the biological process under investigation. However, having access to detailed TFBS information is essential to link a DH site or footprint to a specific TF binding event at the DNA level. Although these experimental methods have substantially increased our knowledge of GRNs in *Arabidopsis*, a systematic profiling of all *Arabidopsis* TFs using ChIP-Seq is currently not feasible (11). Similarly, DH assays still have limitations, as they depend on prior knowledge of TFBSs, the identification of TF footprints requires a good sequencing depth, and DH sites can be condition-specific. Also for TF ChIP condition-specificity is an important factor, as determining protein-DNA binding at different time points can offer new insights in network dynamics or transient regulatory interactions (12,13). New initiatives are being developed that are aimed to profile pools of TFs in a single experiment or study. Two studies used high-throughput *in vitro* assays to identify TFBSs for a large collection of TFs using protein binding microarrays (PBMs) (14,15). Weirauch and co-workers used PBMs to determine the DNA sequence preferences for more than 1000 TFs from 131 species. Although PBM assays can profile TFBSs for different TFs, the detection of BSs is limited to 10-12 nucleotides. The DNA affinity purification (DAP) assay is another *in vitro* method that allowed profiling TFBSs in *Arabidopsis* (16). Out of 1,812 TFs tested, the BSs of 529 TFs were successfully determined, as DAP-seq experiments are affected by different factors like primary sequence, DNA methylation and chromatin accessibility.

Currently there are different databases available which centralize information about plant promoters and TF binding in the promoter regions of target genes. AGRIS, PLACE and PlantCARE integrate collections of consensus BS for TFs from literature (17–19). AthaMAP contains more detailed TF binding specificities modeled as position weight matrices (20). Several tools have been developed that query these databases and identify the TFBSs for a promoter of interest. For example, Athena utilizes the consensus TFBSs for 150 TFs to allow rapid identification and visualization of regulatory elements for a promoter of interest (21). PlantPAN extends this straightforward promoter analysis by performing gene group analysis and predicts the co-occurrence of TFBSs in the promoter (22). CressInt is a web resource which integrates a wide range of gene regulation datasets, including TFBSs, genome-wide histone modifications and open chromatin information (23). As such it does not only identify TFBSs for a promoter of interest but is also able to predict which genetic variants impact the binding ability of a specific TF. Although the aforementioned tools generate a map of TFBSs for plant promoter sequences of interest, many of them use simple mapping of TFBSs to promoters. This approach is highly prone to false positives since TFBSs are often short and typically contain some level of degeneracy (24,25). Furthermore, most tools only integrate a subset of the available TFBS information or restrict their analysis to an arbitrarily defined 500 or 1000bp proximal promoter.

To overcome some of the limitations inherent to experimental techniques, computational approaches making use of publicly available TFBS information can also be used to delineate GRNs. AtRegNet is a tool that allows users to visualize complex networks formed by TFs and their target genes by integrating regulatory interactions from published data (19). Given the limitations associated to simply mapping TFBS to promoters, more advanced filtering schemes have been implemented which increase the specificity to computationally map functional TFBS and construct GRNs. Examples of such filtering steps comprise exploiting conservation of non-coding DNA or the integrating co-regulatory gene information with TFBSs (23,25–31). These filtering approaches, however, do not fully resolve the problem of false positives and might also suffer from false negatives, where functional binding events remain undetected. To tackle some of these challenges, we present TF2Network, a tool that identifies candidate TF regulators for set of co-expressed or functionally related genes based on enriched TFBS. Apart from validating TF2Network using different gold standard datasets, we demonstrate how it can be used to predict new regulators for different biological processes, offering a versatile tool for the improved functional and regulatory annotation of Arabidopsis genes.

## MATERIALS AND METHODS

### Collection of Position Weight Matrices

The motif collection used for TF2Network consists of 1,793 *Arabidopsis* position weight matrices (PWMs), representing 916 TFs from different sources including CisBP (14), Franco-Zorrilla et al. (2014), Plant Cistrome Database (16), JASPAR 2016 (32), UNIPROBE (33), AGRIS (19) and AthaMAP (20). The collected BSs were either in the form of PWMs or consensus sequences and all were converted into position count matrices scaled to 100. TFs were assigned to gene families based on the PlnTFDB 3.0 database (34).

### Extraction of gene regions and PWM mapping

We defined three regulatory gene region types based on the TAIR10 release of *Arabidopsis* and relative to the translation start (TrSS) and end site (TrES) of the gene (35). Long (5000bp upstream from TrSS and 1000bp downstream from TrES), Intermediate (1000bp upstream and 500bp downstream) and Short (500bp upstream i.e. core promoter). For Long and Intermediate, introns were retained while extracting the regions. If another gene is present upstream of the gene, the region is cut where this upstream gene starts or ends. Here, upstream and downstream are used relative to the TrSS and TrES, respectively, because it has been shown that the regulatory elements can be found in the 5’ and 3’ untranslated region (UTR) (4,36–38). Coding regions were not included as it has been shown that the TF binding in coding regions is mostly passive and could also be nonfunctional in the regulation of gene expression (39). The second reason to include UTRs is that not all genes have information about their UTRs.

All PWMs were mapped to the three extracted gene regions using Cluster-Buster (“–c” parameter set to zero) to allow a score for every region without trimming to retain the family specific information within the BS (40). PWM matches mapping inside exons were discarded. If a gene region has more than one match for a specific PWM, only the match with the maximum score was considered for ranking. Ranking of the different gene regions was done based on the mapping score. The PWM gene feature file was created by selecting the top 1000, 2000 or 5000, 7000 and 10000 scoring genes for every PWM. By default the top 5000 scoring genes are used.

### Enrichment statistics calculations

Motif enrichment for given input gene set is calculated using the hypergeometric distribution based on the set of motifs mapped on the gene regions using the probability mass function:

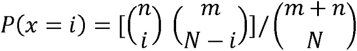

where,

n = #genes with a match for the tested PWM

m = #genes without the PWM match in the genome (having at least one match with the motif collection)

N = #genes in the input set

i = #genes in the input set with the PWM match.

For each enriched motif, the q-value of enrichment is determined using the Benjamini-Hochberg (B&H) correction for multiple hypotheses testing. To empirically evaluate the false discovery rate (FDR), control datasets were generated for every TF by randomly selecting 500 genes from the pool of non-bound ChIP genes. The default q-value is set to 0.05 which, based on 24,000 control datasets corresponds to a FDR of 0.34% (0.17 false positive TFs per run / 50 predicted TFs per run), indicating that the B&H correction effectively controls for false positives.

### ChIP-Seq and TF perturbation gene set benchmarks

In total 24 TFs were selected to build a benchmark containing TF ChIP bound regions (details given in Supplementary Table S1). For 20 TFs peaks were collected from (4) where peaks were annotated to the closest gene in TAIR10 (35). For E2Fa, KAN1 and CCA1 the annotated peaks were downloaded from the original paper (see Supplementary Table S1). For ABI5, the narrow peaks were collected from the Plant Cistrome Database and these peaks were also annotated to the closest gene in the TAIR10 release (16). For WRKY33, as the peaks were not reported in the paper, the raw data was processed. Raw reads for 2 replicates were downloaded from the Sequence Read Archive (SRA) (see Supplementary Table S1) and mapped to the *Arabidopsis* genome using Bowtie2 and only the reads that map uniquely to the genome were considered for peak calling (41). MACS2 was used to identify the peaks for both the replicates and only peaks that had 50% intersection in both of the replicates were considered for the annotation, as only retaining the intersecting region of replicates might miss the actual binding event (42). The reference sample for the intersection analysis was determined as the replicate that had maximum number of peaks. For all TFs, the genes associated with the 500 top scoring peaks were considered for the benchmark gene sets (called ‘ChIP genes’). For all evaluation experiments using the ‘ChIP genes’ benchmark, PWMs derived from the ChIP-Seq experiments were not included in order to avoid circular reasoning.

TF perturbation differential gene expression datasets were obtained for 23 TFs from different publications listed in Supplementary Table S2. Genes were sorted based on the p-value/Fold change (FC)/log(FC) and the top 500 genes were selected for validation. Note that for some TFs less than 500 DE genes were available. If for one TF more than one condition or time point was profiled, each gene set was considered independently for running the TF ‘DE genes’ benchmark. For TFs with multiple DE gene sets, the TF was scored as a true positive prediction when for at least one DE gene set the correct regulator was found. The ‘ChIP genes’ and ‘DE genes’ gold standards are available in Supplementary Datasets S1 and S2, respectively.

### Tool comparison

The ‘ChIP genes’ were used to compare the performance of TF2Network with Cistome and PlantRegMap (43,44). For the Cistome tool, the FIMO-mapped Weirauch Set was used to predict motifs using the option “only significantly enriched motifs” in 1000bp upstream region from the translation start site (no larger promoter sizes can be selected). For calculating ranks, the predicted motifs were sorted by decreasing z-score. Due to the limitation of this tool to take only 50 genes at a time as input, top scoring 50 genes for each TF were submitted (running time approximately 5 minutes). To control the False Discovery Rate (FDR) of PlantRegMap, we used a p-value cutoff of 0.007 as this setting yielded no significant results on the control datasets. This threshold was used to have a fair comparison with TF2Network considering similar levels of FDR.

### Experimental interactions

Experimental interactions included in the web interface were derived from different experimental data sources. For protein-DNA interactions, peaks of all TFs used in the ChIP benchmark along with other TFs from Heyndrickx and co-workers and 21 TF mock samples from Song and co-workers were collected (4,45). These peaks were annotated to the closest gene in TAIR10 (35). The Y1H derived interactions were directly downloaded from the original publications (5–7,46–49). Protein-protein interaction data were retrieved from the studies listed in Table 3 and Gene Ontology (GO) annotations were downloaded from TAIR on 9 May 2017 (35).

Co-expression edges were calculated based on a compendium containing all samples from (50). For each sample accession, all available runs were downloaded as FASTQ files from the SRA (51) and concatenated. Expression quantification was performed with Kallisto (v0.43.0) (52), which produced TPM values for each transcript in the AtRTD2 reference transcriptome (53). Gene-level expression was obtained by summing the TPM values for each gene. Co-expression between gene pairs was calculated as Pearson’s correlation coefficients that were transformed to z-scores for each gene. Only co-expression edges that have a z-score with an absolute value larger than 3 were retained. Co-expression scores for predicted regulators in the webtool are based on these edges with the z-score cutoff.

To evaluate of TF2Network is capable of predicting cooperative TFs, we determined, per input gene set, how many of the top 50 predicted TFs have a known PPI. This procedure was applied for each gene set present in the ‘ChIP genes’ benchmark (n=24). To assess the significance of the observed number of interacting TFs for these real gene sets, controls were generated. For controls, the same procedure was followed as for the real top 50 predicted TFs except that 50 TFs were randomly selected from the database of 916 TFs (10,000 times). The PPI count was defined as the number of TF interactions that were experimentally confirmed by PPIs (source data reported in Table 3). The final p-value was calculated as the number of times the PPI count of a control dataset was greater than that of the real dataset, divided by 10,000.

### Construction and analysis of functional regulons

To construct the functional regulons, only experimental GO annotations were considered (evidence codes EXP, IMP, IDA, IPI, IGI, and IEP). For each gene the top 300 most co-expressed genes were retained and used as input for GO enrichment analysis, in each of the 13 expression atlases present in CORNET (54). GO enrichment analysis was performed using the hypergeometric distribution with q-value cutoff of 1e-5. If a gene was annotated with a certain GO term that was also significantly enriched in its co-expressed genes, this gene set was retained as a valid functional regulon which was subsequently submitted to TF2Network. Out of all predictions for all functional regulons annotated with a specific GO biological process (BP) term, the top 50 regulators were retained. These top regulators were further sorted based on number of times they are predicted in the functional regulons. The most frequently occurring top 50 regulators from this sorted list were annotated as TFs regulating that particular GO biological process. Like this, TFs were identified for functional regulons for all GO BPs considering all 13 expression atlases.

For the unknown Brassicaceae genes, detailed expression information was obtained from Vaneechoutte et.al. 2017 (55). Gene family and phylostrata data was retrieved from the PLAZA 3.0 Dicots comparative genomics database (56). PLAZA 3.0 gene homology data was also used to divide genes into five phylostrata (Viridiplantae, Embryophyta, Angiosperms, Eudicots, and Brassicaceae). GO BP terms used to identify new stress-related TFs are GO:0009408 response to heat, GO:0009409 response to cold, GO:0009651 response to salt stress, GO:0009644 response to high light intensity and GO:0009607 response to biotic stimulus. Expression profiles of genes predicted to control circadian rhythm were evaluated using the Diurnal Tool (<u>http://diurnal.mocklerlab.org/</u>) (57).

## RESULTS

### Basic features of TF2Network

TF2Network takes as input a set of co-regulated, co-expressed or functionally related *Arabidopsis* genes and predicts TF regulators based on enrichment analysis using known TFBSs. In contrast to simple mapping tools that report all TFBS matching to a given set of promoters, efficient enrichment statistics are applied to only return predicted TFBS and regulators for which the corresponding TFBS occurs much more than expected by chance in the input gene set. In total 1,793 motifs corresponding to TFBS for 916 Arabidopsis TFs have been integrated from different databases and literature sources (see Material and Methods). The length of TFBSs varied from 4 to 30 nucleotides (nt) with a median length of 10nt (Supplementary Figure S1 panel A). The length variation was mainly due to the presence or absence of flanking bases surrounding the core BS. For e.g. for TF AT4G25490 (CBF1, which has binding specificity for CCGAC), the BS from CisBP was 10nt long containing the core CCGAC and flanking nucleotides (yrCCGACata). The core of CBF1 BS from DAP-seq was more conserved compared to the BS from CisBP and the length was 15nt (kyyrCCGACatm). All integrated TFBSs cover TFs from 62 different TF families and the number of TFs for each family is shown in Supplementary Figure S1, panel B, along with the distribution of lengths of BSs within each family.

For predicted regulators selected by the user, an interactive GRN visualization was developed that makes it possible to explore in more detail the regulatory interactions for a specific TF or a set of target genes. Furthermore, experimental protein-protein and protein-DNA interaction data, co-expression information as well as gene function information were integrated in order to improve the extraction of biological information from the predicted networks. For every gene, a pre-defined non-coding DNA region is defined which covers the exon-masked gene body, the full upstream (up to 5kb) and the 1kb downstream sequence, which is used to first map the motif collection and subsequently to calculate the TF enrichment for a given input gene set. The primary output of TF2Network comprises a list of regulators and the associated target genes, together with the enrichment statistics for each motif and information about co-expression between the input genes and the predicted regulator.

### Evaluation of TF2Network using different benchmarks

To evaluate the performance of TF2Network, two experimental benchmark datasets were constructed. The first benchmark is based on TF ChIP bound genes (called ‘ChIP genes’) for 24 different TFs while the second dataset contains differentially expressed genes after TF perturbation (called ‘DE genes’) for 23 TFs. These datasets were collected from different studies that performed TF ChIP binding assays and TF perturbation experiments (more details in Materials and Methods; Supplementary Table S1 and S2). We evaluated TF2Network on three gene region types, long, intermediate and short, extracted from the Arabidopsis genome (5kb upstream + exon-masked gene body + 1kb downstream, 1kb upstream + exon-masked gene body + 500bp downstream, and 500bp upstream core promoter, respectively). The performance of TF2Network was evaluated by measuring how many times it correctly recovered the TF from the ChIP or DE gene sets and scoring at which rank the correct TF was reported. For the ‘ChIP genes’, the overall performance of TF2Network varied between 75-92% (Figure 1A). Out of the three different gene regions, the best performance was observed for the long region with 92% recovery (22/24 TFs) compared to 83% and 75% for intermediate and short, respectively. Despite the potential risk of having a higher level of false positive TF predictions due to a larger search space, the advantage of using the long region type is that it captures TF binding events that are beyond the 1kb upstream promoter. For KAN1 and WUS, this type of distal TFBS covers 39% and 28% of all binding events, respectively. Considering all 24 TF ChIP gene sets, on an average 31% of the TFBSs are missed by both the intermediate and short gene regions. Furthermore, 4% of all binding events that lie in the introns are missed by the short region. For the 22 correctly predicted regulators using the long gene regions, 62% of the TFs were correctly identified within the top 5 predictions (called top 5 ranks). Apart from the TF level recovery, the performance of TF2Network was also evaluated at the TF family level (Figure 1A and 1B). To evaluate the family level performance, we counted how many times TF2Network found a regulator from the same family as the TF profiled in the benchmark datasets. Overall, the family level performance ranged from 83% to 96% and was again highest (23/24) for the long gene region type. The performance of the tool was also evaluated for top 200 ‘ChIP genes’, yielding very similar results as for the top 500 ‘ChIP genes’ (Supplementary Figure S2). Based on these results, the long gene region is used by default in the TF2Network tool.

**Figure 1:**
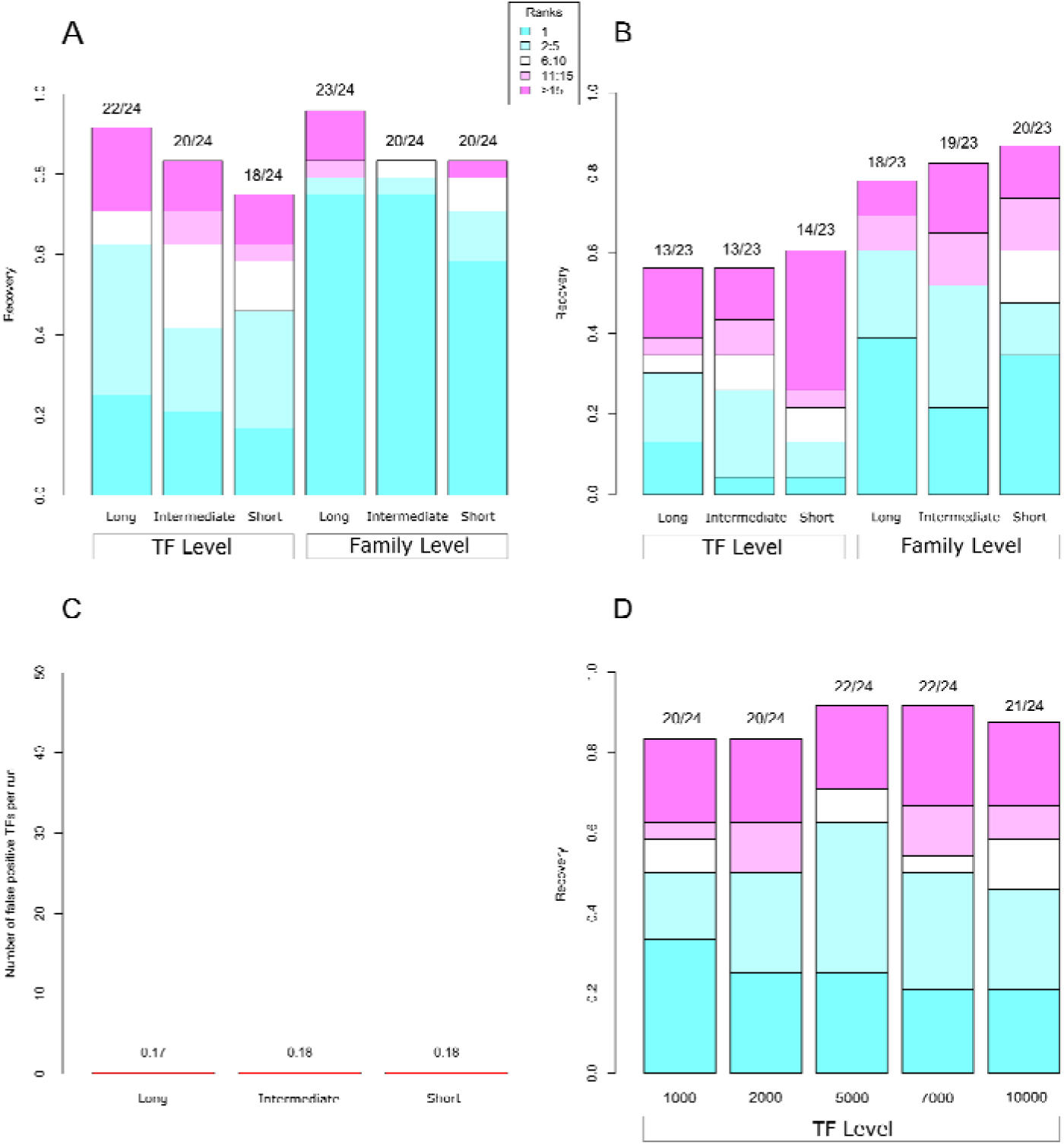
TF2Network benchmarking for ChIP bound, DE and Control gene sets. Recovery on Y-axis corresponds to the fraction of TFs correctly predicted within the benchmark. (**A)** The performance of TF2Network for ‘ChIP genes’ (N = 24 TFs). The colors of stacked bars represent the rank bins with colors from cyan to magenta indicating rank 1, rank 2-5, rank 6-10, rank 11-15 and rank 15 onwards. (**B)** The performance for ‘DE genes’ consisting of 23 TFs. (**C)** The performance of TF2Network for control gene sets based on genes that were not bound in a ChIP experiment. (**D)** Feature comparison for the ‘ChIP genes’ benchmark. Top 1000, 2000, 5000, 7000 and 10000 ranked genes per TF were evaluated.

‘DE genes’ collected from different TF perturbation experiments are genes that transcriptionally respond to either overexpression or knockdown of a specific TF (Supplementary Table S2). In these benchmark datasets, there is no information if the perturbed TF is binding to and directly regulates the DE genes or not. Figure 1B shows the overall performance of TF2Network for the ‘DE genes’. Although the overall performance is identical for long and intermediate regions (56%), the long region has better TF level recovery within top 5 ranks for both TF level (30%) and at the TF family level (60%) recovery compared to the other two regions (48% and 52% for short and intermediate region, respectively). This shows that TF2Network is also able to predict the correct regulator for gene sets regulated by a specific TF including both direct and indirect target genes. At the TF family level, the performance ranges between 75 and 87% for the short and long gene regions, respectively.

### Assessing the robustness of TF2Network

Apart from analyzing the performance of TF2Network on the two gold standard benchmarks, we also assessed the specificity of the method using control gene sets (Figure 1C). To measure false positive predictions, for each TF in the ChIP benchmark dataset a control gene set was constructed by randomly selecting 500 genes from the pool of non-bound ChIP genes. When running TF2Network on these control gene sets with the default q-value threshold of 0.05, on an average 0.17 TFs were predicted per control gene set (long gene regions), corresponding to an FDR of 0.34% (see Material and Methods). The FDR for predicting the TFs for the other gene regions was highly similar.

As TF2Network performs motif enrichment analysis using a PWM gene feature file, we also verified if the number of selected top-scoring PWM gene matches has an influence on the overall performance (see Material and Methods). In total, we selected seven PWM gene feature files, which contain the best 1000, 2000, 5000, 7000 and 10000 gene matches, respectively. The performance at the TF level for the different PWM feature files is shown in Figure 1D. Feature 5000 had an overall recovery of 22/24 correct TFs which was better than using the 1000 and 2000 PWM gene feature files. Although the PWM gene feature files containing 7000 matches had same overall recovery, the 5000 feature file shows the best performance when considering the top 5 and top 10 ranks. Therefore, the PWM gene feature file containing the best 5000 matches per PWM is used as the default.

Although the two benchmarks used to evaluate TF2Network are strongly enriched for true target genes for a specific TF, it is important to verify how robust the TF predictions are for input gene sets with lower signal-to-noise ratios. Therefore, we generated additional datasets where starting from the ‘ChIP genes’ benchmark containing 500 bound regions per TF, increasing fractions of non-bound targets were included (denoted as percentage of noise in Supplementary Figure S3). Whereas the original recovery of 92% (22/24) correct TF predictions drops to 79% and 63% when adding 30% and 50% noise, respectively, the fraction of correct predictions in the top 5 ranks only drops from 62% to 54% when adding 50% noise. These results indicate that the performance of TF2Network is robust even in the case where the TF signal in the input gene set is low.

### Comparison of TF2Network with existing plant TF analysis tools

The performance of TF2Network was compared with previously published tools that also aim to predict BSs or TFs for a set of input genes. Although many tools offer PWM mapping on a set of input genes, only Cistome and the TF enrichment tool of PlantRegMap return overrepresented BSs or TFs in their output and thus were selected to compare with TF2Network (43,44). Like TF2Network, both tools take as an input a set of genes for which the users wants to predict regulators. Here we used the ‘ChIP genes’ to compare the performance of the different tools. The overall recovery for Cistome was 21% for these 24 TFs and this tool does not allow to directly link the enriched PWM to the associated TF. Cistome incorporates a small collection of TFBSs and also runs slow for the settings used for this comparison (see Tool comparison in Materials and Methods).

PlantRegMap uses a Fisher’s exact test to calculate TF enrichment using TF binding site data combined with DNase I hypersensitive sites, TF footprints and histone modifications. PlantRegMap shows an overall performance of 67% (16/24, detailed ranks in Table 1), which is lower than TF2Network. As PlantRegMap uses 500bp upstream and 100bp downstream from the transcription start site, it will miss the target genes where the TF binding is more upstream or downstream. Considering all tested TFs present in the ‘ChIP genes’ benchmark, no cases were observed where PlantRegMap or Cistome predicted the correct TF but TF2Network failed to do so. However, there is some complementarity in the predicted ranks, as for TFs like BES1, EIN3, FLC, the ranks of PlantRegMap are better than the ranks predicted by TF2Network.

**Table 1:**
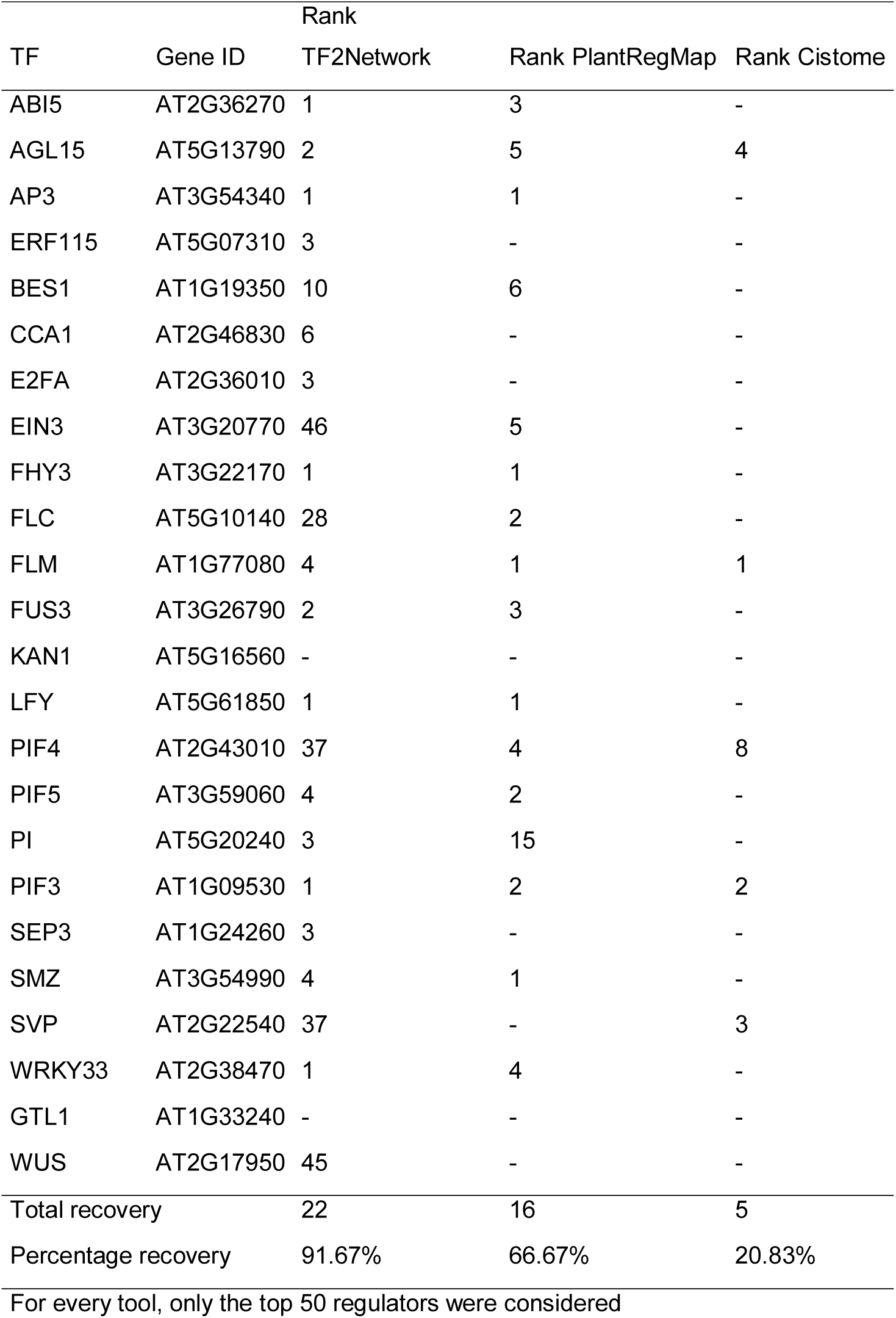
Performance comparison different tools

### Validation of target genes of TF regulators predicted by TF2Network

Apart from evaluating the performance of TF2Network to correctly predict regulators for the different benchmark gene sets, we also assessed if the predicted target genes for a given regulator would correspond with functional targets. Through the combination of TF perturbation gene expression information and ChIP-Seq data, we quantified how well predicted target genes based on DE input genes overlapped with ChIP-bound genes. Selecting four TFs for which a correct regulator was predicted within the top five predictions (PIF4, FHY3, EIN3 and PIF3; see Table 2), the average recovery of target genes by TF2Network confirmed by ChIP was 69%. An overview of validated target genes for PIF3, FHY3 and PIF4 are shown in Figure 2. In case the correct TF was not predicted in the top 10 predictions, the recovery was lower, ranging from 25 to 52%. Overall, these results show that TF2Network is not only predicting a correct regulator in many cases, but also succeeds to link a regulator to its downstream target genes with good sensitivity.

**Table 2:** Validation of target genes predicted by TF2Network

| TF Symbol | Gene ID | Rank from TF2Network | #Target<br>Genes | #TG confirmed by<br>ChIP | Percentage<br>Recovery |
| --- | --- | --- | --- | --- | --- |
| PIF4 | AT2G43010 | 1 | 45 | 25 | 55.56% |
| FHY3 | AT3G22170 | 1 | 84 | 79 | 94.05% |
| EIN3 | AT3G20770 | 1 | 106 | 65 | 61.32% |
| PIF3 | AT1G09530 | 5 | 43 | 29 | 67.44% |
| PIF5 | AT3G59060 | 13 | 43 | 15 | 34.88% |
| AP3 | AT3G54340 | 23 | 102 | 53 | 51.96% |
| PI | AT5G20240 | 32 | 103 | 50 | 48.54% |
| BES1 | AT1G19350 | 36 | 40 | 10 | 25.00% |

**Table 3:** Overview experimental regulatory interactions included in TF2Network

| Interaction type | Number of interactions | Comment <sup>(1)</sup> |
| --- | --- | --- |
| Protein-DNA interactions | 167,636 | ChIP-Seq: 53 TFs |
|  | 3,247 | Y1H: 652 TFs |
| Protein-protein interactions | 76,236 |  |
| Co-expression | 13,139,852 | Correlation transformed into z-score and cutoff of z-score>3 or z-score <3 was used |

<sup>(1)</sup> For a full overview of all included published studies, see Supplementary Table S5

**Figure 2:**
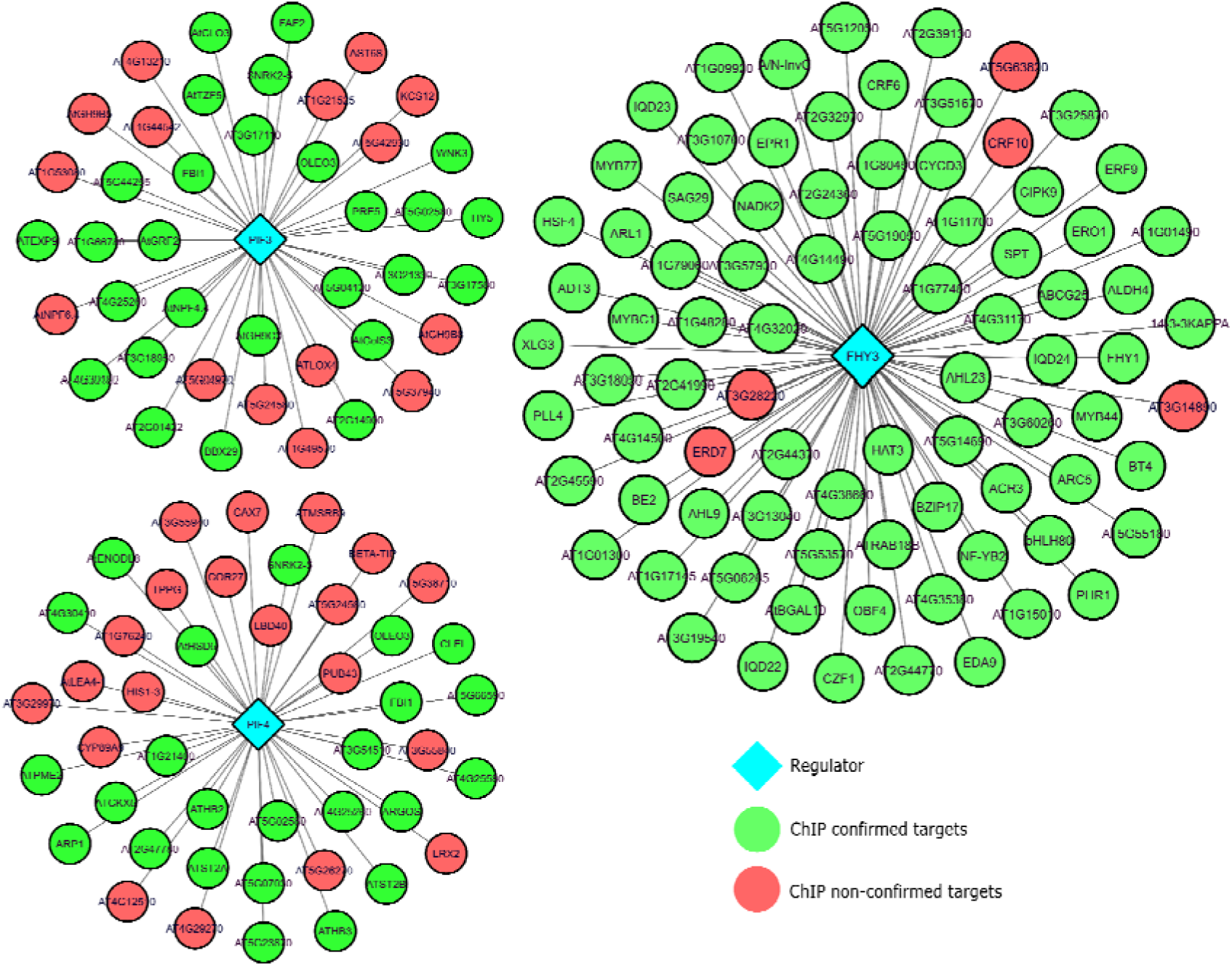
Validation of TF2Network target gene prediction. The network figure shows the results for a set of PIF3, PIF4 and FHY DE input genes. Green nodes denote target genes predicted by TF2Network that are confirmed by ChIP experiments while red genes were non-confirmed target genes.

### Protein-protein Interactions between predicted TFs reveal cooperative TFs

Co-binding TFs achieve increased DNA-binding specificity through the formation of higher-order protein complexes, which can modify the DNA-binding affinity of the individual TFs in the complex. Depending on whether the cooperative TFs are from the same or from different TF families, we here call these interactions homotypic or heterotypic, respectively. To evaluate if TF2Network can shed light on putative cooperative TFs, we used the regulator prediction results of the 24 ‘ChIP genes’ benchmark datasets to identify TFs physically interacting with each other through a known protein-protein interaction (PPI). Specifically, for the top 50 predicted regulators per TF gene set, the number of interacting TFs was counted and compared with control datasets to assess the significance of the observed overlap between the predicted TFs and the PPI data (see Material and Methods). The distribution of PPI counts for real and control gene sets is shown in Figure 3A and reveals that for 13/24 TF gene sets a significant number of interacting TFs can be found (based on a p-value< 0.01 threshold, dashed line Figure 3A). We further verified if these combinations of predicted TFs offer new insights on homotypic versus heterotypic TF interactions. Figure 3B shows the number of homo and heterotypic interactions for the 13 TF sets deemed significant in panel 3A. Overall, these results indicate that TF2Network can detect both type of interactions (63% and 37% homo and heterotypic interactions, respectively). For the FLM ‘ChIP genes’ genes, 147 experimentally validated PPIs were found, of which 98 and 49 were homo- and heterotypic interactions, respectively. Among the heterotypic interactions detected for these 24 TF gene sets, many of them have previously been described including interactions between MYB-bHLH, TCP–MYB–bHLH, BES1–bHLH, bHLH–bZIP, AP2/EREBP–HB etc. (58). Figure 3C shows a detailed overview of the predicted TFs and associated PPIs for the FLM ChIP gene set. This network view offers a better understanding of predicted TFs, which were clustered based on family information, and their interactions with different TFs from the same or a different TF family, such as MADS, TCP, BBR/BPC, bHLH, C2H2 and Pseudo ARR-B. Known interaction for FLM comprise PPIs with AG, AGL31, AGL68-70 (MADS family) and TCP14 (TEOSINTE BRANCHED, cycloidea and PCF family). In conclusion, these results indicate that among the predicted regulators returned by TF2Network frequently cooperative TFs can be found, facilitating the generation of new hypothesis about TFs operating through protein complexes to control specific sets of spatial-temporal co-regulated genes.

**Figure 3:**
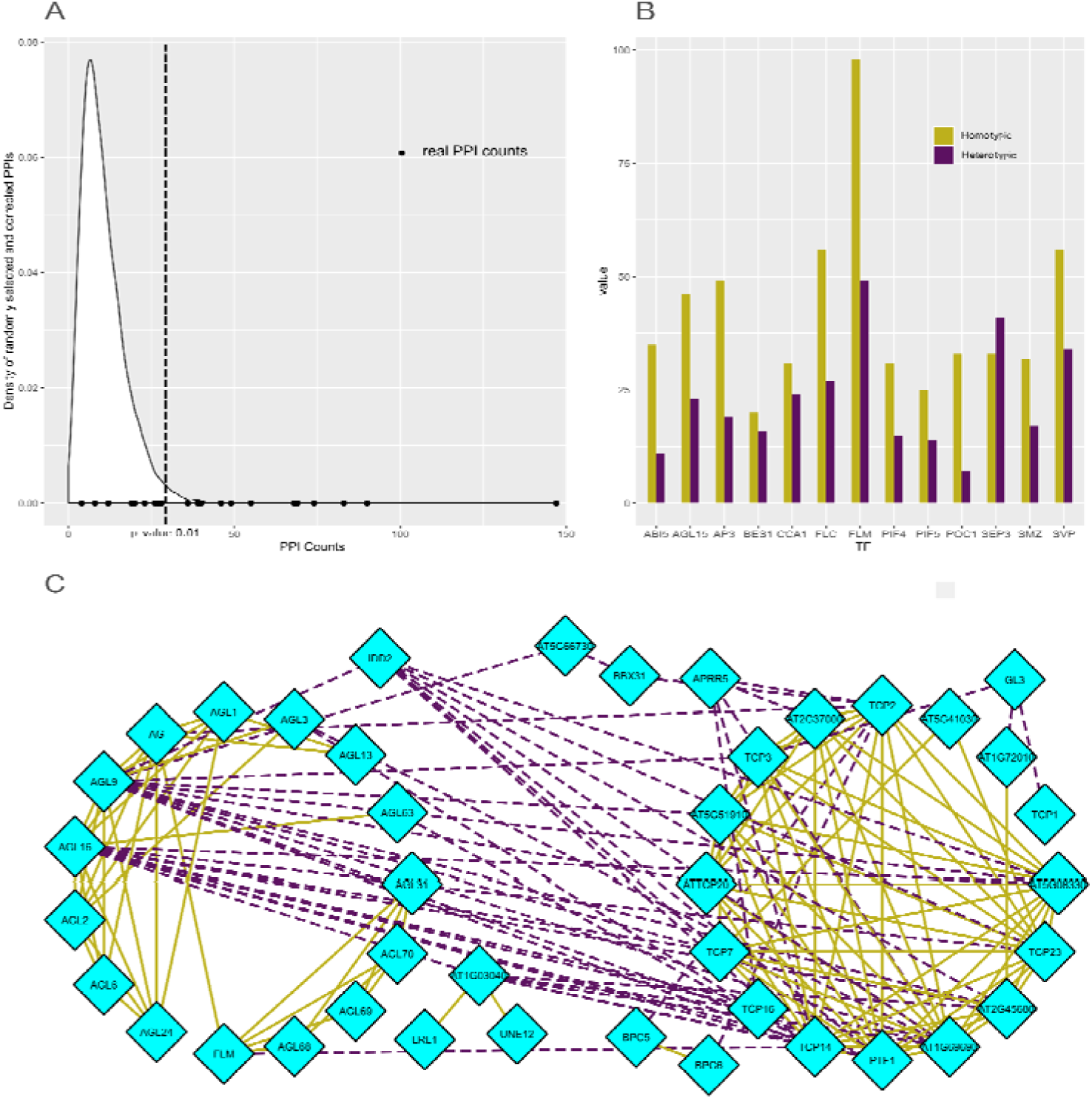
Significance testing of PPI interactions in TF2Network output. **(A)**Density distribution of experimental PPIs within sets of 50 randomly selected TFs are shown in white (n=10,000 control sets). Black dots represent real counts for PPIs within top 50 regulators predicted by TF2Network for 24 ‘ChIP genes’ sets. **(B)** Distribution of homotypic and heterotypic PPIs for sets with p-value < 0.01 in panel A are shown in green and purple, respectively. **(C)** PPI network view for regulators of FLM showing homotypic and heterotypic interactions within TFs covering different TF families.

### TF2Network web interface

To make the TF2Network algorithm easily accessible and to allow intuitive exploration of the predicted GRN, an online interface was created (see Availability for URL). On the start page, the user can provide a set of genes as a list of gene identifiers or aliases. Submitting the gene set will start the retrieval of PWM motif mappings, gene co-expression data, protein-DNA interactions and protein-protein interactions (PPI), and Gene Ontology (GO) annotations from the server (see Material and Methods), after which the TF2Network algorithm is run client-side. TF2Network performs hypergeometric tests on motifs in the input genes and ranks these according to the resulting q-values. These ranks are used to order TFs in the regulator panel (Figure 4A). This panel shows all predicted TFs, the percentage of input genes with which they show Z-score co-expression (CO), the percentage of input genes for which the predicted regulator has known experimental protein-DNA interactions (PD) and a list of their PWMs that are significantly enriched in the input gene set. Z-score co-expression reports the fraction of target genes that are strongly positive or negatively correlated with the predicted TF (see Material and Methods). As shown for the ‘ChIP genes’ benchmark in Supplementary Figure S4, high Z-score co-expression frequently coincides with correct TF predictions. For each PWM the rank, q-value, and number of input genes that contain the PWM are shown. Hovering over the PWM in the table shows the source of this PWM along with its consensus sequence information displayed as a sequence logo. All prediction results can be exported to CSV format by clicking the ‘Export predictions’ button at the bottom of the regulator panel.

**Figure 4:**
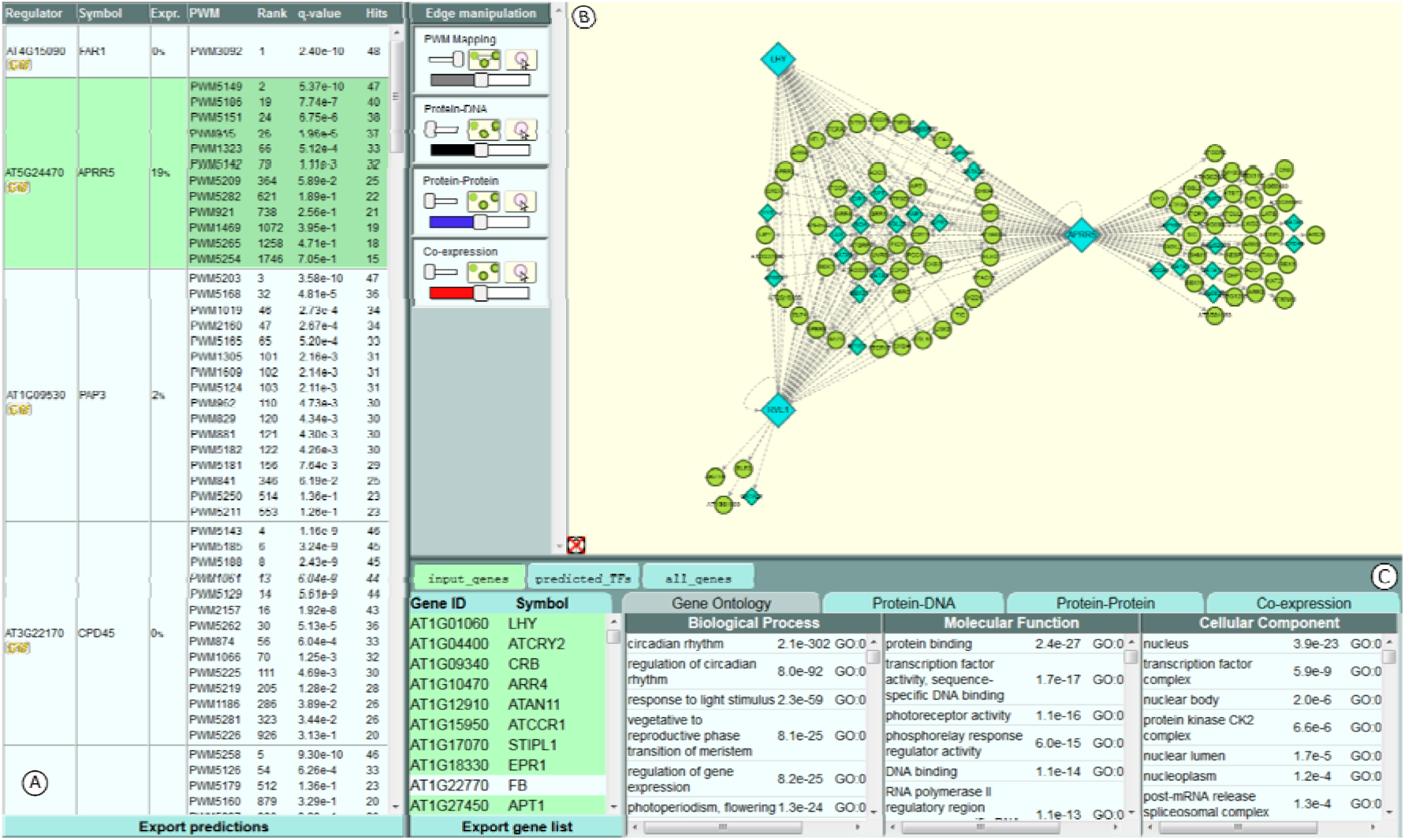
Screenshot of the TF2Network user interface. **(A)** The regulator panel listing predicted regulators. **(B)** Cytoscape panel showing GRN where blue diamonds represent TFs and green nodes refer to non-TF genes. **(C)** Gene info panel that shows gene ontology enrichments on gene sets

Selecting PWMs in the regulator table starts a network visualization of the corresponding TFs and the input genes that contain the selected PWMs in the Cytoscape panel (Figure 4B). Multiple TFs can be active in the visualization at a time. The positions of the gene nodes in the visualization reflects the number of active TFs they are predicted to be regulated by: genes that contain selected PWMs for all active TFs are placed centrally in a compact circle. The remaining genes with PWMs for more than one active TFs are then placed in concentric circles around the center so that genes with more TFs are closer to the center. Next, the active TFs are spaced out evenly outside these concentric circles. Finally, genes with selected PWMs for only one of the active TFs are placed in a compact circle near the TF, distal to the center (Supplementary Figure S5, panel A).

PWM Mappings, Co-expression, PPIs, and Protein-DNA interactions are shown as grey, red, blue, and black edges between genes in the network, respectively (Supplementary Figure S5, panel B and C). How and when each edge type is shown can be adjusted in the Edge manipulation boxes on the left in the Cytoscape panel. For each edge type, the box contains one large slider to change the thickness of the edges and another small slider to toggle between hiding all edges, showing only edges between selected genes, and showing all edges of this type. A button is also provided to enable the display of all edges for a gene when hovering the mouse over it. For PWM Mappings, only edges for currently selected PWMs in the regulator panel are shown. A full overview of all edge types and how to manipulate the different edges is explained in a Tutorial present on the TF2Network website.

When a single gene is selected in the Cytoscape panel, additional information will be displayed in the gene info panel below (Figure 4C). The Gene Ontology tab will contain all GO terms associated to the selected gene. Three additional tabs are provided that contain tables of genes with which the selected gene has a Protein-DNA, Protein-Protein, or Co-expression interaction. Hovering over or clicking the headers of these tables will highlight or select all interacting genes for this edge type. The PWM Mapping tab shows a visualization of the selected gene’s structure and the positions of the active PWMs on the genes introns, upstream, and downstream regions. The visualization can be dragged to pan horizontally and zoomed in or out by scrolling. Zooming in closely also shows the genomic DNA sequence (Supplementary Figure S5, panels D and E). Multiple genes can be selected by holding the SHIFT key while clicking genes or dragging the mouse. The gene info panel will then show a prompt to save the selection as a new gene set. When a gene set is saved, the Gene Ontology tab will display all GO terms for genes in the set, sorted by their enrichment p-value. Hovering over or clicking a GO term will highlight or select the associated genes in the set. The other three tabs will provide a button to download the corresponding interactions between genes in the set and a button to reposition the genes in the Cytoscape panel so that the distance between interacting genes is minimized. Saved gene sets will be shown at the top of the gene info panel. When TF2Network is initialized, three gene sets will already be available: (i) the input set provided by the user (‘input_genes’), (ii) a gene set containing all predicted regulators (‘predicted_TFs’), and (iii) a gene set containing the union of the first two sets (‘all_genes’). Within the web tool, hovering the mouse over certain buttons, sliders, tabs, or panels will show a tooltip describing its functionality. Tooltips can be disabled by clicking the icon in the bottom left corner of the Cytoscape panel.

### Systematic regulatory annotation of biological processes

Based on the good performance of TF2Network in the different benchmark experiments, we next asked if the tool can be used to perform systematic regulatory annotation of TFs and target genes. Although in Arabidopsis 1,441 TFs have GO annotations based on experimental support, still 751 TF lack experimental functional annotations. As for >150 of these unknown TFs high-quality BS information is available, TF2Network can be used to predict which genes and biological processes they regulate. Furthermore, based on the latest GO annotations, still 11,447 Arabidopsis protein-coding genes lack any functional data. Prior to generating functional annotations for unknown genes, we first executed a large-scale analysis to measure how well TF2Network can recover known TF functional annotations.

To perform a systematic functional analysis of TFs in Arabidopsis, we first combined GO Biological Process (BP) annotations with co-expression information to identify functional regulons. A functional regulon is a set of co-expressing genes which can be annotated to a specific biological process, using GO functional enrichment analysis (here only considering experimental GO annotations, see Material and Methods). In order to make use of the large number of transcript profiling studies performed for Arabidopsis and to capture condition-specific co-regulatory information, functional regulons were delineated in 13 different microarray expression atlases covering a general expression compendium, stress, biotic and abiotic stress, AtGenExpress, development, flower, leaf, root, seed, whole plant, genetic modifications and hormone covering thousands of available Affymetrix arrays (54). Starting from experimental annotations for 1,918 different GO BP terms and considering the general expression compendium, 20,548 genes could be assigned to one or more functional regulon (in total 2,367 regulons). For regulons annotated with a specific GO term, we applied TF2Network to verify if known TF regulators could be recovered. For example, starting from the GO term circadian rhythm (GO:0007623), 36 functional regulons could be identified. After running TF2Network on these 36 regulons and retaining the top 50 predicted TFs with the highest ranks, 9 regulators known to control circadian rhythm were successfully identified. Examples of correctly predicted TFs are RVE2 (AT5G37260), HY5 (AT5G11260), RVE7 (AT5G52660), CCA1 (AT2G46830) and LHY1 (AT1G01060). Examples of other processes for which many known TFs were recovered are defense response (15 TFs), plant organ development (12 TFs), hormone-mediated signaling pathway (8 TFs) and cell differentiation (8 TFs).

Considering all functional regulons in all 13 expression atlases, we could identify known functional annotations for 471 TFs, from 34 different TF families, covering 500 different GO BP terms. A known functional annotation refers to a TF predicted to be a regulator for a functional regulon annotated with specific BP term, and for which experimental GO data is available supporting the role of that TF in that BP. Interestingly, we observed that for a given BP term, the number of correct regulators sometimes varied strongly depending on the expression atlases that was used to define the regulons (Supplementary Table S3). Regulation of post-embryonic development (GO:0048580) is an example where in the development and flower atlases 12 TFs were correctly predicted, respectively, while in the other atlases only 0-4 correct TFs were recovered. Examples of development related TFs uniquely detected in the development atlas include AT1G33240 (GTL1), AT2G22540 (AGL22), AT5G10140 (AGL25) and AT3G15270 (SPL5). This result confirms that using condition-specific expression data improves the inference of gene functions (59,60). Apart from recovering known TF functions in one specific expression atlas, we also observed cases of strong complementarity. For root system development (GO:0022622), per expression atlas between 0-5 correct regulators were identified, while in total 9 TFs known to be involved in root development were recovered. A full overview of all correctly predicted TFs for all GO BP terms is given in Supplementary Table S3.

All functional regulons with known TFs assigned to it offer an interesting dataset to systematically characterize unknown Arabidopsis genes, as these regulons contain experimentally characterized genes assigned to a specific GO BP term as well as TFs known to regulate these processes. For the 471 correct TFs identified in functional regulons in one of the 13 expression atlases, regulatory and functional interactions were found for 19,609 target genes. Considering the genes not annotated to any biological process using experimental or computational GO annotations, 6,498 of the 11,447 unknown protein-coding genes were successfully annotated to one or more BP. As unknown Arabidopsis genes are strongly enriched for young gene families (61), we focused on a set of genes uniquely found in the genus Arabidopsis or the Brassicaceae family (called Brassicaceae phylostratum). Based on gene family information from the PLAZA 3.0 Dicots database, we identified 2,253 unknown genes belonging to the Brassicaceae phylostratum of which 591 were assigned to functional regulons with known regulators. Interesting, 116 of these genes were part of regulons annotated with flower development, which formed a GRN of 288 regulatory interactions including 29 known TFs involved in flower development (such as AGL15, FLM, SEP3, FLC and PI). Forty-three of these genes were predicted to be regulated by three or more flowering TFs and were analyzed in more detail. Overlapping these genes with detailed RNA-Seq based tissue expression information revealed that 18 genes were expressed in specific stages of flower development. In addition, for four genes evidence for TF binding based on experimental protein-DNA interactions was found (covering AGL15 and PISTILLATA TFs; see Supplementary Table S4). For a gene family present in all Brassicaceae species encoding proteins with a domain of unknown function DUF1216 (family HOM03D007139), three out of five genes were part of flower development regulons and found to be strongly expressed in anthers (Supplementary Table S4). Two other genes, AT1G05540 and AT1G28375, part of families HOM03D000799 and HOM03D012901 having homologs in multiple Brassicaceae species, were each found to be expressed in flowers and bound by the flowering TF AGL15. A full overview of all known TFs, their associated GO BP functional regulons and the predicted target genes which are currently unknown can be found in Supplementary Dataset S3.

To validate some of the new predictions associating TFs to specific BPs, we examined some specific expression datasets which were not present in our atlases. A first study performed detailed transcript profiling in multiple stress conditions, covering cold, heat, high-light, salt and biotic stress (62). To be consistent, we here only focused on 5 GO BP core stresses corresponding to the conditions analyzed in this study (see Material and Methods). To validate the TF2Network predictions for these stresses, the predicted TFs for these BPs were compared with the regulators being differentially expressed in one of these five stresses. For every stress, the overlap between predicted TFs and stress-responsive TFs was measured. Figure 5A shows the TFs that are confirmed either by experimental GO annotations or by the expression data of Barah and co-workers. For salt stress, 12 predicted TFs were confirmed of which eight are new TFs showing differential expression in the salt stress expression dataset. Examples of newly predicted TFs are AT3G17609 (HYH) which is known to respond to UV light, AT3G46640 (LUX) which is involved in regulation of circadian rhythm, AT1G32640 (MYC2) which is responsive to jasmonic acid and controls defense response, and AT4G01120 (bZIP54) which responds to the blue light. The WRKY TF family consists of a huge collection of TFs that are regulators of plant immune response (63). Figure 5A shows that 16 of these known WRKY TFs were correctly identified playing a role in biotic stress response. TF2Network adds two more TFs, AT4G23550 (WRKY29) and AT4G01250 (WRKY22), as being regulators of biotic stress. AT1G46264 (HSFB4) and AT2G40350, which are known to be involved in cell division and heat acclimation, respectively, were predicted by TF2Network, and confirmed by the differentially expressed dataset, to be involved in high light stress. In total 22 new TFs predicted to control specific stress responses were confirmed by the stress-specific differentially expressed genes.

**Figure 5:**
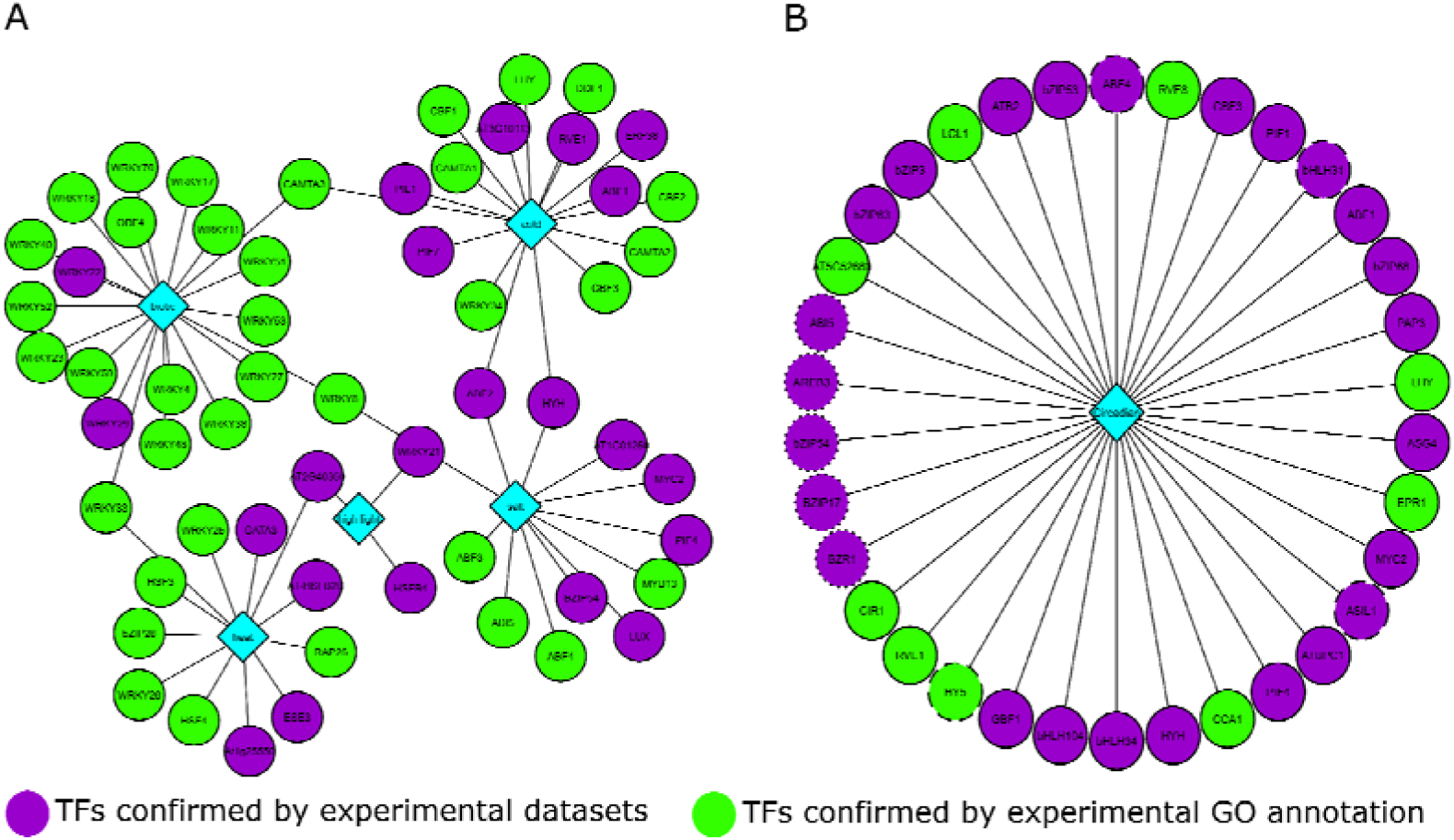
Validation of new TF functions. **(A)** Validation of TFs predicted in different stress conditions using expression data from Barah et al. 2015 for five stress conditions. Diamonds indicate GO BP terms while nodes denote TFs. Purple nodes represent TFs confirmed using experimental GO annotations and green nodes represent TFs validated using the stress-specific expression data. **(B)** Validation of TFs predicted to be involved in circadian rhythm. Green nodes had a correlation >= 0.8 reported by the diurnal tool for longday or shortday conditions. Dashed and dotted nodes correspond to the longday and shortday condition, respectively. TFs confirmed in both the conditions are shown using solid lined nodes.

In a second evaluation, TFs predicted for circadian rhythm were evaluated using the Diurnal Tool (57), which describes the circadian genome-wide expression of genes measured using microarrays. For the GO term circadian rhythm, the top 50 TF predictions for general atlas were submitted to the Diurnal tool to examine if they show a circadian expression pattern. Furthermore, known circadian rhythm TFs based on experimental GO annotations were also scored. For longday conditions, 29/50 TFs showed a diurnal expression pattern, out of which 9 were newly predicted TFs and 20 were TFs previously described to be involved in circadian rhythm (Figure 5B). In shortday conditions 30/50 of the predicted TFs were confirmed to show a diurnal pattern, 8 of which were known TFs involved in circadian rhythm. The detailed circadian rhythm expression patterns are shown in Supplementary Figure S6 for longday and shortday, respectively.

## DISCUSSION

In this study, we leveraged publicly available TFBS information to computationally identify candidate TFs regulating sets of co-expressing or functionally related genes. Validation of TF2Network using a gold standard dataset based on experimental TF binding data revealed that it recovers 92% of the true regulators using the long region promoter definition. In addition, processing negative control gene sets and gene sets with a decreasing number of true TF-bound target genes showed that TF2Network controls false positives well and is robust to noise in the input gene sets. These results demonstrate that the combination of PWM mapping combined with applying a stringent enrichment test offers a good trade-off between obtaining the correct regulators and controlling for false positives. The good performance of TF2Network on the long region definition has the advantage that potential TF binding events in introns or downstream of the gene are also identified. In the ‘ChIP genes’ benchmark covering *in vitro* binding data for 24 TFs, on average 31% of all binding events were located outside the proximal 1000bp promoter. Thus, in contrast to a recent analysis which concluded that a majority of TFBSs (86%) lie in the region from −1,000□bp to +200□bp with respect to the transcription start site (64), our benchmark indicates that many regulatory interactions are missed by promoter analysis tools only considering a 1000bp upstream promoter. Although we also evaluated other filter schemes based on open chromatin information or using conserved non-coding sequences (9,10,25,26), these approaches had a lower recovery of correct regulators in our benchmarks. This finding is in agreement with a recent study that identified putative *cis*-regulatory elements in high salinity stress conditions, where a reduced sensitivity was observed when applying filtering using open chromatin or conserved non-coding sequences (65). In contrast to TF-bound genes measured by ChIP, ‘DE genes’ offer a more realistic but also more challenging dataset to evaluate TF2Network, as this benchmark also contains genes that are indirectly regulated by the perturbed TF (66). The overall recovery of 56% of the correct regulators indicates that TF2Network also works well for input gene sets where not all genes are directly regulated by a specific TF. The good performance of TF2Network at the TF family level reveals that the tool can also be used to identify candidate TF families controlling a specific biological process. This family-based approach offers a practical means to identify candidate regulators in cases where only one or a few TFs of a specific TF family have detailed binding site information.

By combining the TFs predicted by TF2Network on the ‘DE genes’ benchmark with the experimental TF binding data from different ChIP experiments, the recovery of functional target genes was evaluated. In case the correct regulator was predicted in the top 5 predictions, on average 69% of the predicted target genes were confirmed by TF binding measured through ChIP. Obviously, as both ‘ChIP genes’ and ‘DE genes’ identified through transcript profiling after TF perturbation are obtained from *in vivo* samples which might cover different conditions, obtaining a perfect overlap between both profiling methods is unrealistic. In addition, very frequently ChIP binding events lack a consensus TFBS, which could mark indirect or non-functional binding events. For example, for six TFs involved in flowering the overlap between ‘DE genes’ and TF bound genes measured using ChIP varied between 7-22% (67). Therefore, the good recovery of bound and regulated target genes through the integration of co-expression information with detailed BS information in TF2Network confirms that, even in the absence of DE genes after TF perturbation, functional target genes can successfully be recovered (4). Compared with other publicly available tools to study TF gene regulation in plants, TF2Network offers several advantages. Firstly, it integrates the largest number of TFBSs currently available for Arabidopsis. Secondly, the performance of TF2Network to predict correct regulators is better compared to Cistome and PlantRegMap. For large input gene sets covering hundreds of genes, the running time of TF2Network is between 5-10 seconds while identifying significantly enriched motifs for 50 genes using Cistome requires around 5 minutes. In contrast to TF2Network, both Cistome and the TF enrichment tool in PlantRegMap lack a network view where enriched BSs are connected to candidate target genes. Thirdly, through a newly developed web interface the user can interactively and efficiently delineate a GRN for a set of input genes. Advanced network layout visualizations were implemented making it possible to simultaneously study multiple predicted TFs for an input gene set, generating new insights on the complexity of TF control. In addition, experimental regulatory datasets from >15 different studies were included, making it possible to efficiently integrate >76,000 known protein-protein and >170,000 known protein-DNA interactions while exploring the predicted GRNs. Several download options are also available to study the predicted regulators, target genes or the associated networks using other tools. Fourthly, we demonstrated how combining the predicted regulators with protein-protein interaction data makes it possible to recover known co-binding TFs, both for TFs from the same or different families, and to generate new insights about cooperative TFs operating via protein complexes.

Through the integration of experimental GO BP annotations and co-expression information, functional regulons were delineated as a starting point to perform a systematic functional and regulatory analysis of Arabidopsis genes. For 471 TFs, known functional annotations were correctly predicted by TF2Network for a wide range of BPs, indicating that functional regulons based on experimental GO annotations offer an interesting dataset to link TFs to functionally coherent sets of target genes covering both known and unknown genes. Starting from 6,498 unknown genes part of functional regulons with correctly identified regulators, we characterized a set of 43 genes only found in Arabidopsis or Brassicaceae species involved in flower development that were predicted to be complexly regulated by 3 or more flowering TFs. Sixty-five percent of these genes were confirmed to be expressed in specific flower developmental stages or to be bound by flower TFs measured in ChIP experiments for well-studied flowering regulators. Furthermore, we identified 25 new TFs predicted to be involved in circadian rhythm, which were confirmed by strong diurnal expression profiles. In addition, 22 newly predicted TFs controlling various types of (a)biotic stress were confirmed using differential gene expression data from Barah and co-workers (62).

In conclusion, we have shown that known TF functions and functional regulatory interactions can successfully be identified when using differentially expressed genes or functional regulons as input for TF2Network. This result reveals that context-specific expression data together with detailed TFBS information has great potential to enlarge our understanding of gene regulation in Arabidopsis. Furthermore, through the integration of protein-protein interactions as well as co-expression information, it offers an invaluable resource to study combinatorial TF control in more detail in plants.

## AVAILABILITY

The TF2Network tool is available at <u>http://bioinformatics.psb.ugent.be/webtools/TF2Network/</u>

## SUPPLEMENTARY DATA

- Supplementary Figure S1. Distribution of lengths and family information for PWM collection.
- Supplementary Figure S2. Performance of TF2Network for ChIP200 and ChIP500 genes.
- Supplementary Figure S3. Performance of TF2Network after adding non-bound genes.
- Supplementary Figure S4. Overview of Z-score co-expression values for the ‘ChIP genes’ benchmark.
- Supplementary Figure S5. Overview of available network layouts and integrated experimental data starting from the Demo dataset
- Supplementary Figure S6. Diurnal pattern of genes in longday and shortday conditions.
- Supplementary Table S1. Overview of TF ChIP datasets used for benchmarking
- Supplementary Table S2. Overview of TF DE gene sets for benchmarking
- Supplementary Table S3. Overview of correct TF-GO functions based on functional regulons in different expression atlases
- Supplementary Table S4. Overview of unknown genes part of the Brassicaceae phylostratum annotated with flower development
- Supplementary Table S5. Overview of studies included in Table 3
- Supplementary Dataset S1. ChIP genes
- Supplementary Dataset S2. DE genes
- Supplementary Dataset S3. Functional and regulatory annotations for unknown Arabidopsis genes. This table lists per GO BP term (columns 1 and 2) the known TF (column 3) and the unknown target gene (column 4) predicted to be involved in this BP. Note that per target gene multiple times the same GO BP term can be present because of different predicted regulators.

## ACKNOWLEDGEMENT

We thank Leelavati Narlikar for proofreading the manuscript, Hana Imrichova, Gert Hulselmans and Stein Aerts for their technical assistance.

